# Discovery of the first small-molecule extracellular inhibitor of K_Ca_3.1

**DOI:** 10.64898/2026.03.08.710400

**Authors:** Joana Massa, Jurek Hense, Tanja Gangnus, Matteo Gozzi, Emma Etmar Bulk, Bjoern B. Burckhardt, Martina Düfer, Albrecht Schwab, Oliver Koch

## Abstract

The ion channel K_Ca_3.1 plays a role in immune regulation, red blood cell function, and is linked to numerous types of cancer. Various animal toxins, such as maurotoxin, bind to the extracellular side of K_Ca_3.1, providing a potential starting point for inhibitor development. We report in this work the discovery of a novel, small-molecule inhibitor, with a micromolar IC_50_, which was specifically designed to target plasma-membrane K_Ca_3.1 channels from the extracellular side. This compound can serve as a starting point for the development of more selective inhibitors and probes.

For the identification of new extracellular inhibitors, molecular dynamics simulations were performed using the experimental structures of K_Ca_3.1 and maurotoxin. The simulations produced a validated binding mode, highlighting key residues involved in the interaction between the toxin and the channel. These findings laid the foundation for the structure-based identification of novel extracellular small-molecule inhibitors of K_Ca_3.1. The Molport database, containing approximately 50 million compounds, was screened using protein-ligand docking, yielding a hit molecule that was experimentally confirmed using patch clamp assays.

## Introduction

Ion channels represent important drug targets and fall into two main classes based on their gating mechanisms: ligand-gated ion channels (LGICs) and voltage-gated ion channels (VGICs). These classes also differ structurally; while LGICs have large extracellular domains that often feature important ligand binding sites, VGICs typically lack this feature.(Alexander et al., 2023; Huang et al., 2024) VGICs consist of two to four subunits each containing a voltage-sensing domain (VSD) with the four helices S1 to S4 and two additional helices (S5, S6) that form the channel pore.(Huang et al., 2024)

The intermediate-conductance calcium-activated potassium channel 3.1 (K_Ca_3.1) shares this typical architecture of VGICs but remains insensitive to voltage changes. It lacks positively charged residues in its helix S4 which would be important for voltage sensitivity.(Catterall, 2010; Cassell et al., 2025) Instead, K_Ca_3.1 is activated by intracellular Ca^2+^. Upon Ca^2+^ binding, Calmodulin (CaM) binds to the intracellular side of the channel, which opens the channel pore and enables K^+^ flux. The pore, located at the channel centre, is lined by all four channel subunits. The narrowest point of the pore is formed by V282, the hydrophobic gate.(Lee and MacKinnon, 2018; Ong et al., 2025)

K_Ca_3.1 regulates immune cell function and is associated with multiple cancer types (Wulff and Castle, 2010; Wulff and Köhler, 2013; Brown et al., 2018; Luca Matteo Todesca, 2021) like non-small cell lung cancer where K_Ca_3.1 drives progression. Blocking K_Ca_3.1 enhances treatment effectiveness of the EGFR tyrosine kinase inhibitor erlotinib in non-small cell lung cancer cells.(Todesca et al., 2024) K_Ca_3.1 also regulates red blood cell (RBC) size making it a suitable target for the treatment of sickle cells disease, a genetic disease impairing RBC function.(Xiao, 2024; Ong et al., 2025) K_Ca_3.1 channel blockers were developed that bind in the pore below the selectivity filter directly blocking ion flow. Senicapoc, one such inhibitor, reached Phase III clinical trials, but failed to meet the primary end point of pain reduction despite improving RBC clinical parameters.(Oikonomopoulou et al., 2021; Lee et al., 2024)

Most importantly, K_Ca_3.1 is not only found in the plasma membrane, but it was also found in mitochondria. Expression of the channel in the inner mitochondrial membrane is linked to cancer cell migration and progression.(Bachmann et al., 2022; Bulk et al., 2022) Small molecules like senicapoc cross the membrane and inhibit both plasma membrane and mitochondrial channels. Furthermore, senicapoc only inhibits the open-state as this channel conformation is required to reach its binding site.(Thale et al., 2025) Extracellular block offers the advantage that it is state-independent compared with pore blockers. Therefore, extracellular blockers that do not cross the membrane would be a valuable chemical probe for a better understanding of both channel types with the potential to be developed into a new type of drugs.

Here, we present an exhaustive virtual screening campaign including high-throughput docking of 50 million molecules that led to the first known extracellular small molecule blocker of the K_Ca_3.1 channel. The modelled binding mode of known peptide toxins like maurotoxin were used as a starting point for the structure-based screening campaign. These toxins are specifically targeting the plasma membrane channel as their size and polarity prevent membrane crossing. Unfortunately, these toxins lack K_Ca_3.1 specificity, limiting their usefulness as tool compounds. Charybdotoxin (ChTx) binds to K_Ca_3.1 with a K_d_ of 5 nM but shows similar affinity for K_v_1.2 (K_d_ = 14 nM). Another toxin called maurotoxin (MTx), extracted from the venom of the scorpioid *Scorpio maurus*, has the highest affinity for K_Ca_3.1 with a K_d_ of 1 nM but similarly blocks K_v_ channels with low nanomolar affinity.(Blanc et al., 1997; Rochat et al., 1998; Chandy et al., 2004) For MTx the toxin residues Lys23 and Tyr32 were found to be essential for its pore-blocking ability via mutagenesis.(Castle et al., 2003) Efforts to enhance K_Ca_3.1 selectivity over voltage-gated potassium channels failed to abolish K_v_ channel activity completely.(Castle et al., 2003) Additionally, these toxins are less readily available and synthetically accessible than small molecules. This highlights the urgent need for a selective extracellular blocker.

## Methods

In the following, channel residues will be indicated using one-letter code, while toxin residues will make use of the three-letter code for amino acids to more easily distinguish the peptide and channel.

### Structure Preparation

For K_Ca_3.1, the cryo-EM structure of the open-pore channel with the PDB ID 6CNO(Lee and MacKinnon, 2018) was used as a template. For MTx, the NMR structure with the PDB ID 1TXM was used.(Blanc et al., 1997) All protein structures were separately loaded into MOE (Vilar et al., 2008) version 2022. For the channel, missing residues of all four transmembrane subunits and CaM were modelled. For the bound CaM residues M1, A2 and K149 were added using the “Protein Builder”. The missing channel loop L124 - Q141 was built using the “Loop Builder”. The C-terminal channel residues S387 - K427 were taken from the crystal structure with the PDB ID 6D42.(Ji et al., 2018) All protein structures were then prepared using the “Quick Prep” tool. This routine included the protonation at a pH of 7.4 with “Protonate 3D” and energy minimisation using the AMBER10:EHT forcefield at a root-mean-square gradient of 0.1 kcal/mol/Å. The final structures were visually inspected to check structural integrity.

### Protein-Peptide Docking

The prepared protein structures were used to model the toxin binding. First, a flexible protein-peptide docking was done employing the HADDOCK webserver version 2.4.(Honorato et al., 2024) The prepared structures were uploaded, and ambiguous interaction restraints (AIRs) were set to include pre-existing knowledge about the putative toxin/channel interaction interface. As active residues of MTx, Lys23 and Tyr32 were chosen, which have been previously shown to be essential for toxin binding to the channel.(Castle et al., 2003) As active residues of the channel G254, D255, V256, V257, Q229, A230, A233 and T234 were defined, which are all located on the extracellular side of the channel. G254 and D255 form the extracellular part of the ion conductance pore, while V256 and V257 are two residues directly surrounding the extracellular side of the pore. Q229, A230, A233 and T234 belong to the turret region, which is important for toxin binding.(Chen and Chung, 2015) Otherwise, default settings were chosen and the results obtained through the server were downloaded. The results were clustered by HADDOCK using the Fraction of Common Contacts with a 60% cut-off and a minimum cluster size of four. The docking pose chosen for further refinement was selected for its favourable energy terms, the best Z-score, the HADDOCK score, and the highest buried surface area (**Figure 3A**).

### Molecular Dynamics Simulations of the Channel-Toxin System

The selected docking pose of MTx to K_Ca_3.1 was used as the starting structure for the simulations. Using CHARMM-GUI (Wu et al., 2014; Lee et al., 2016) a 160 x 160 Å^2^ 1-palmitoyl-2-oleoyl-sn-glycero-3-phosphocholine (POPC) bilayer was constructed. Then the channel-peptide complex was inserted into the lipid bilayer using the InflateGRO (Kandt et al., 2007) script, which led to the deletion of overlapping lipids and a system with dimensions of 170 x 170 x 220 Å^3^. The final structure consists of 615 POPC molecules. The channel and toxin parameters were taken from the AMBER99SB-ILDN force field (Lindorff-Larsen et al., 2010). For the lipids, the Slipids2016 (Klauda et al., 2010; Jämbeck and Lyubartsev, 2012) forcefield and to model water TIP3P (Jorgensen et al., 1983) was used. As the open, activated structure was used, which features four CaMs attached to the channel, the system includes 12 Ca^2+^ as each CaM subunit has three Ca^2+^ bound. Moreover, three K^+^ ions were modelled bound to the channel pore to stabilise the selectivity filter region in the simulation. This was suggested by instabilities in the pore region, observed in preliminary MD simulations without these ions (**Figure S1**). The coordinates of the three K^+^ bound in the ion conductance pore were taken from the overlayed closed structure (PDB ID 6CNM) to the channel-peptide complex.(Lee and MacKinnon, 2018) Further K^+^ and Cl^-^ were added in a concentration of 150 mM to neutralise the system. Electrostatic interactions were calculated using Particle-Mesh-Ewald.(Darden et al., 1993) First, the system was energy minimised using the steepest descent algorithm for 50,000 steps with position restraints on all heavy atoms. All simulations use a time step of 2 fs. This was followed by three separate equilibration simulations.

First, the system was heated to 310 K using the V-rescale thermostat and equilibrated for 5 ns. Then, applying pressure of 1 bar via the Berendsen barostat, the system was further equilibrated for 1 ns while heating the system to 323 K using the Berendsen thermostat. As a final equilibration step, the system was simulated for 10 ns using the Parinello-Rahman barostat to maintain a pressure of 1 bar. Semi-isotropic pressure scaling was applied. As a thermostat, V-rescale was used and the temperature was kept at 323 K. The settings of the last equilibration step were kept for the production simulations. In all simulations, the cut-off for short-range van-der-Waals and electrostatic interactions was set to 1.2 nm. Furthermore, LINCS constraints were always applied to H-bonds.(Hess et al., 1997) During equilibration, restraints were applied on all protein and lipid heavy atoms as well as the ions bound to the protein. For the production simulations, all restraints were removed and coordinates were output every 100 ps. System preparation and production simulations were performed using GROMACS 2020.6.(Abraham et al., 2015) In the production simulations a peak performance of around 20 ns/day was reached using two RTX2080 Super GPUs and two Intel Xeon Gold 5220 CPUs. The toxin-channel system was simulated for at least 500 ns in five independent repeats, accumulating in a total simulation time of 2.785 µs (**Table S1**). 2785 of the concatenated simulation frames were analysed in the following steps. The average structures of the most populated clusters aligned on the Cα atoms of the channel were visualised and the corresponding RMSD of the channel and peptide was analysed using GROMACS. Further analysis was carried out using custom Python scripts utilising MDAnalysis (Gowers et al., 2016).

### Preparation of the Molport Database

The Molport database was downloaded in SMILES format in June 2023 via the vendor’s ftp protocol. At the time of download, the Molport database contained around 48 Mio. molecules. First the downloaded SMILES strings were protonated at a pH of 7.4 using the OpenBabel(O’Boyle et al., 2011) command line tool. If multiple tautomer were possible only the dominant form was kept. Then, one low energy conformer per compound was constructed using OpenBabel. The prepared coordinates were saved as SD Files and used as input for the docking-based virtual screening campaign.

### Docking-based Virtual Screening Campaign

GOLD (Jones et al., 1997; Verdonk et al., 2003) was used for all docking steps. More details about the docking are shown in **Figure 1** and **Table S1**. Based on the important interactions between MTx and the channel, constraints were defined to accelerate the docking process (**Figure 3F**). The binding site was defined as 15 Å around a central point slightly above the ion channel pore, based on the position of the nitrogen from Lys23.

**Figure 1.**
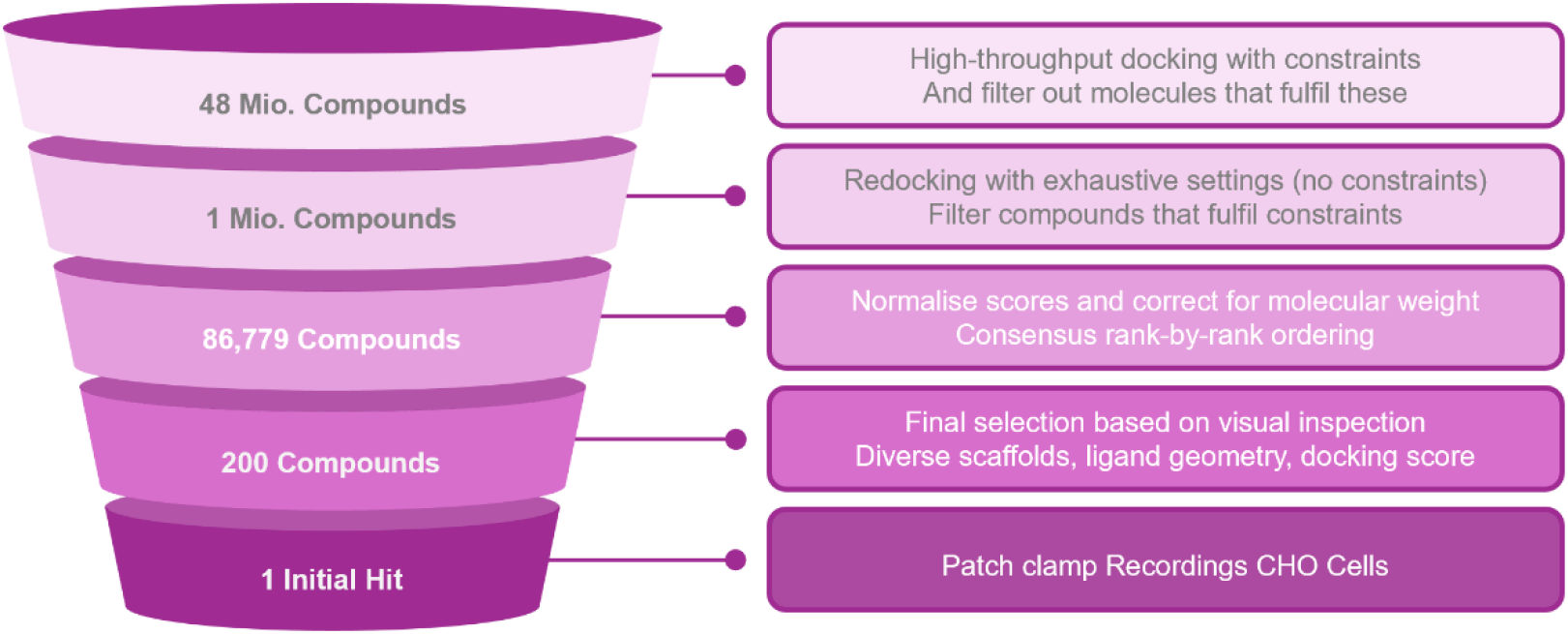
Graphical depiction of the workflow of docking-based virtual screening campaign.

The docking protocol followed a scheme developed earlier.(Koch et al., 2013) The side chain of D255 of subunit D was set to “Rotate” in all docking steps. The rotamer angles were taken from the cryo-EM structure and the GOLD rotamer library. In brief, the protocol consists of three steps: 1) Docking with constraints, 2) more exhaustive free docking and 3) rescoring (**Table S1**). The final scores of all four scoring functions were normalised and corrected for molecular weight. Negative scores resulting from noncompliance with the constraints were set to 0. Molecules for synthesis were selected based on a consensus rank-by-rank scheme (Oda et al., 2006) after the correction step. The consensus score is defined as the average of the individual rankings for all four scoring functions (**Figure 1**). After this first, score-based step, all 15 poses of the 200 highest consensually ranked molecules were visually inspected. Finally, 26 molecules were chosen for synthesis. Analysis of molecular properties was additionally conducted with MOE (Vilar et al., 2008) and Datawarrior (Sander et al., 2015) version 06.01.00.

### Compound synthesis

Of the 26 molecules from the first selection, 21 were synthesised by Enamine Ltd. (Kiev, Ukraine) while five were bought from Molport, SIA (Riga, Latvia). A second quantity of the first hit molecule (**1**) was then purchased from Enamine, as well as ten molecules (**2**-**11**) that expand the structure of **1**. The structures of the first selection of 26 molecules are shown in **Figure S2** and the structure of the hit series, as well as the reference compound senicapoc (**12**), can be seen in **Figure 2**.

**Figure 2.**
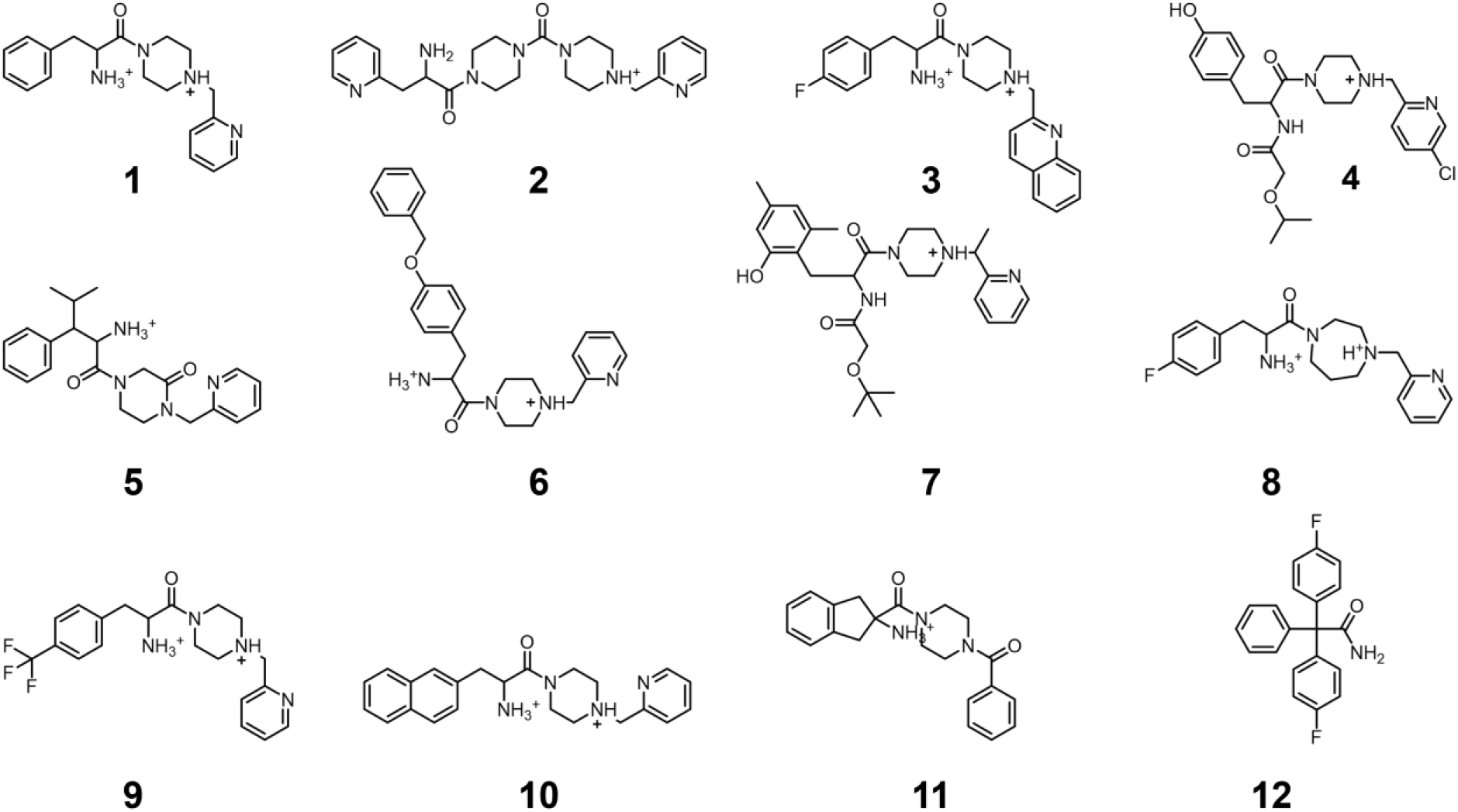
**1** Hit molecule. **2-11** Molecules expanding the series. **12** Senicapoc, molecule serving as positive control in experimental validation.

**Figure 3.**
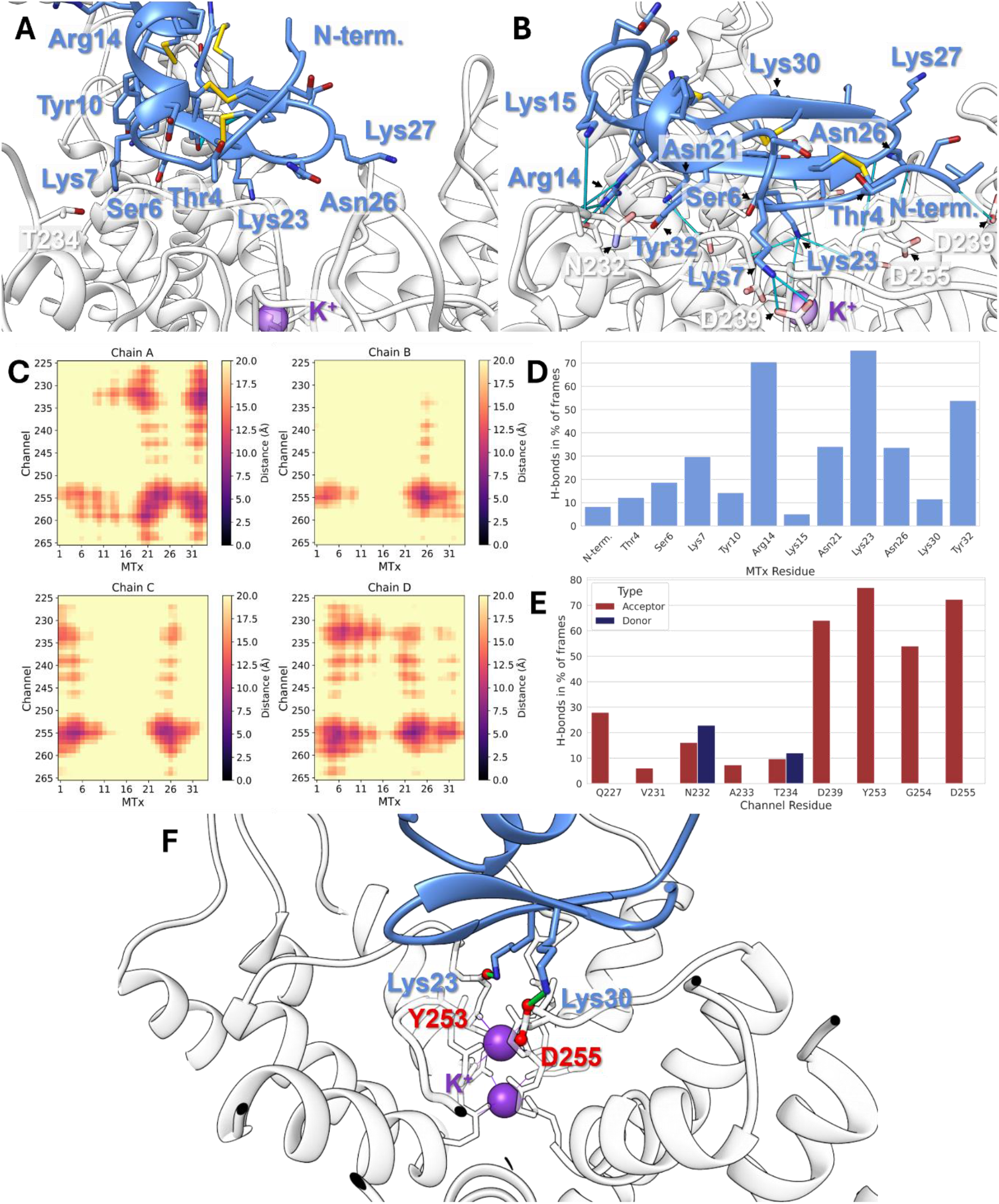
Results of MD simulation of KCa3.1 together with Maurotoxin. In all subfigures the channel is coloured white and the toxin is coloured blue. The protein backbone is shown as ribbons while important residues are displayed as sticks. H-bond interactions are visualised as light blue (**A, B, G**) or green (**F**) sticks. Oxygen atoms are shown in red and nitrogen atoms are coloured dark blue. **A**) Channel-peptide docking pose obtained through the HADDOCK web server.(Honorato et al., 2024) This was used as the starting frame for the MD simulations. **B**) Depiction of the most representative frame of the largest cluster identified from MD simulation of MTx bound to K_Ca_3.1. **C**) The four plots illustrate the average Cα atom distance between all MTx residues (x-axis) and each channel subunit (y-axis, only pore and turret region shown). The darker the color the lower the average distance in the MD simulation frames. **D**) MTx residues that are involved in H-bond interactions between the toxin and K_Ca_3.1 **E**) K_Ca_3.1 residues that serve as H-bond donors (blue) and/or H-bond acceptors (red) in interactions between the channel and MTx. **F**) Visualisation of interactions that are defined as constraints in the virtual screening.

### Patch clamp Recordings CHO Cells

Patch clamp recordings with CHO cells were performed by ICE Bioscience (Beijing, China). First, the hit molecule was validated there and afterwards the series extending the hit compound was tested by the same company. All measurements were done as a single repeat except for the IC_50_ determination, which was a double repeat. CHO cells stably expressing K_Ca_3.1 were used in this study. The CHO cells were cultured at 37°C, 5% CO_2_ in F-121X+GlutaMAX medium supplemented with 10% FBS. For patch clamp recordings 8×10^3^ cells were seeded on 12 mm coverslips. Whole-cell patch clamp recordings using a HEKA EPC10 amplifier were performed. The intracellular solution consisted of 120 mM KCl, 10 mM HEPES, 1.75 mM MgCl_2_ × 6 H_2_O, 8.55 mM CaCl_2_ × 2 H_2_O, 4 mM Na_2_ATP, 10 mM EGTA and the extracellular solution consisted of 160 mM NaCl, 4.5 mM KCl, 1 mM MgCl_2_ × 6 H_2_O, 2 mM CaCl_2_ × 2 H_2_O, 10 mM D-glucose and 10 mM HEPES. All compounds were applied extracellularly. Cells were held at a potential of −80 mV before a hyperpolarizing step to −120 mV for 0.02 s was applied. This was followed by a ramp from −120 mV to +40 mV lasting for 0.2 s. After a second hyperpolarization to −120 mV for 0.02 s, cells were held at −80 mV. This protocol was repeated at 10-second intervals to observe the effect of the putative inhibitors on the peak of K_Ca_3.1 current. Cells were superfused with control solution until the current amplitude stabilized. Compounds were applied until the current reached an equilibrium block (around 5 min). After that the next concentration was tested. The control and test solutions flew sequentially through the chamber from low to high concentration via a gravity-fed solution perfusion system. During the experiment, the solutions were withdrawn from the chamber by a peristaltic pump. Within each patch clamp recording, the current values in response to the compound (Peak compound current) were normalised to blank control (Peak control current) and the inhibition rates were calculated ((1 - Peak compound current)/Peak control current). Mean and standard error (SE) were calculated for each test group, and data are presented as mean ± SE.

All tests were performed at room temperature. Individual curves of the extended series patch clamp recordings are depicted in **Figure S3** and exemplary individual traces are shown in **Figure S4**.

### Protein-Ligand Docking of Extended Series

The molecules were prepared by loading the SMILES of the ten compounds (**Figure 2**, 2-**11**) into MOE. The protonation was set at a pH of 6.5 to ensure protonation of at least one amine group. The 3D conformations were reconstructed and minimised under the MMFFx94. In the first docking, the experimental structure of K_Ca_3.1 prepared as previously described was used. In GOLD, the rotation of the side chain of D255 was disabled this time. The binding site was defined as previously described. GA autoscale was set to 200% and 100 poses per ligand were generated. The same settings were used in the second docking of the molecules **2**-**11** to the K_ATP_ channel. Here, the experimental protein structure of human K_ATP_ in complex with the centipede toxin SpTx1 (PDB ID 9KGL) was used and the standard MOE QuickPrep workflow was applied for protein preparation in this case.

### Molecular Dynamics Simulations of Extended Series

For the MD simulations of compound **1**, the workflow described for the channel-toxin system was followed. In brief, InflateGRO (Kandt et al., 2007) was used to insert the channel-**1** complex into a POPC bilayer and then the system was equilibrated following the scheme previously described. Three independent production runs were conducted, each with a simulation time of at least 500 ns. For the simulations of compounds **2**-**11**, the docking poses of the channel-small-molecule complex were prepared for simulation using CharmmGUI (Wu et al., 2014; Lee et al., 2016). The prepared protein structure from above was used. However, the modelled C-terminus was modified by deleting residues S387-K427 to enable the use of a smaller box size and thereby more efficient simulation. The small molecule parameters were taken from CGenFF (Vanommeslaeghe et al., 2010) and Charmm36m (Huang et al., 2016) was used for the protein and lipids. The complexes were inserted into the membrane using PPM2.0, only the transmembrane domains A-D were submitted to the server. Charged C- and N-termini were chosen and a water thickness of 22.5 was used. 150 mM KCl was added to neutralise the system. TIP3P (Jorgensen et al., 1983) was used to describe the water molecules. The box was initially defined as 150 x 150 Å^2^, and the final systems after initial equilibration were around 144 x 144 x 146 Å^3^. The systems contain around 200,000 atoms and for equilibration, the standard CharmmGUI protocol was applied, which is already described here.(Thale et al., 2025) The production simulations were run at 1 bar and 300 K. Three 500 ns production runs were conducted per compound. Analyses were performed as previously described. **Table S2** summarises all conducted simulations.

### Patch clamp recordings of K_slow_ currents

Pancreatic beta cells were isolated from adult male and female C57BL/6N mice (Charles River, Germany and own breeding, Germany) by collagenase digestion of isolated pancreata to obtain islets followed by trypsin-EDTA digestion. Cells were cultured in RPMI-1640 medium supplemented with 10% FCS, 100 U/mL penicillin and 100 µg/mL streptomycin for up to two days. Patch pipettes with a resistance of 3-5 MΩ were obtained by pulling borosilicate glass capillaries. K_slow_ currents were recorded with an EPC-10 patch clamp amplifier (HEKA, Germany) and analysed with Fitmaster v2×92 (HEKA, Germany). K_slow_ currents were elicited by a train of 26 voltage ramps from the holding potential of −40 mV to 0 mV and back to −40 mV within 200 ms each. K_slow_ currents were measured before and 3 min after the application of the test compound. External solution (in mM): 140 NaCl, 3.6 KCl, 2.5 CaCl_2_, 2 NaHCO_3_, 0.5 NaH_2_PO_4_, 0.5 MgSO_4_, 5 HEPES, 15 glucose, pH adjusted to 7.4 with NaOH. Pipette solution (in mM): 10 NaCl, 10 KCl, 70 K_2_SO_4_, 4 MgCl_2_, 2 CaCl_2_, 10 EGTA, 5 HEPES, pH adjusted to 7.2 with KOH. Amphotericin B was added right before the experiment to the pipette solution (0.25 mg/mL).

### Mass spectrometric determination of LogD value and permeability

A customized high-performance liquid chromatography coupled with mass spectrometry (HPLC-MS/MS) method was developed for the quantification and analysis of **1, 9** and **12** (senicapoc). Chromatographic separation was achieved on an Agilent 1200 SL series system (Agilent Technologies, Ratingen, Germany) equipped with a degasser (G1379B), a binary pump SL (G1379B) and a column oven TCC SL (G1316B) using a Luna PFP(2) 100 Å LC column (100 × 2.0 mm, 3 µm; Phenemonex, Aschaffenburg, Germany)). The mobile phase consisted of 0.1% formic acid (FA) in water (A) and 0.1% FA in methanol (B). A gradient elution was performed at a flow rate of 0.3 mL/min under the following conditions: 0.0–2.0 min: 5% B; 2.0–6.5 min: 5–95% B; 6.5–9.0 min: 95% B. The injection volume of 10 μL was applied with a PAL HTC-xt autosampler (CTC Analytics AG, Zwingen, Switzerland) and the column temperature was maintained at 50 °C. Detection was carried out on an API 4000 triple quadrupole mass spectrometer (AB Sciex, Darmstadt, Germany) equipped with an electrospray ionization (ESI) source. The instrument was operated with an ion spray voltage of 5500 V, a source temperature of 500 °C, curtain gas of 40 a.u., collision gas of 8 a.u., gas 1 of 40 a.u. and gas 2 of 60 a.u. The following transitions were monitored in positive ionization mode and multiple reaction monitoring: Compound 1 325.2 →178.2 (declustering potential (DP) 70, entrance potential (EP) 10, collision energy (CE) 28, collision cell exit potential (CXP) 15), Compound 9 393.1 → 178.2 (DP 80, EP 10, CE 32, CXP 15), senicapoc 324.2 → 200.2 (DP 86, EP 10, CE 31, CXP 5), and the reference Propranolol 260.2 →116.2 (DP 87, EP 11, CE 24, CXP 12), and Propranolol d7 267.2 → 189.2 (DP 82, EP 13, CE 25, CXP 14).

### Distribution Coefficient (logD) Determination

The logD determination was based on the shake flask method optimized for mass spectrometric analysis.(Andrés et al., 2015) The distribution of test compounds was investigated between buffer pre-saturated octanol and octanol pre-saturated buffer using an Andrew+ pipetting robot (Waters Corporation, Eschborn, Germany). LogD was determined at pH 7.4 using 0.1 M potassium phosphate buffer, as well as pH 5.0 and pH 2.2 using 0.15 M citric acid-/phosphate buffer, adjusted to the respective pH. A 10 mM solution of the test compounds in DMSO and the control propranolol were diluted 1:100 in buffer. A 50 µL aliquot was directly taken and 1:1 diluted in dimethyl sulfoxide (v/v, standard). To the residual buffer volume, octanol was added in different v/v ratios, depending on the predicted logD.(Andrés et al., 2015) Samples were shaken for one hour to reach equilibrium before the two phases were separated by centrifugation at 13.200 x g for 10 min. Determinations were performed in triplicate for each compound from the aqueous phase after mixing 1:1 (v/v) with DMSO. For the calculation of logD, the buffer/octanol v/v ratio was used in which the distribution of drug was most similar between octanol and buffer in order to obtain the most precise measurement. The logD was then calculated by the following Equation 1:

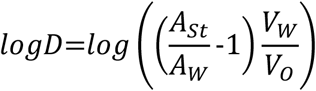

**Equation 1.** Calculation of logD. A_St_: area ratio standard, A_w_: area ratio after partition, V_w_: volume aqueous phase, V_o_: volume octanol

### In vitro Permeability Measurements

The permeability of the substances was determined using Kerski diffusion cells.(Kerski, 2020) Permeation experiments were performed at pH 2.2, 5.0 and 7.4 to investigate the influence of ionization state on membrane permeability. At pH 7.4 0.1 M potassium phosphate buffer was used as donor medium and the acceptor medium consisted of 2% buffered bovine serum albumin in 0.1 M potassium phosphate. At pH 2.2 and 5.0 experiments were performed using a 0.15 M citric acid/phosphate buffer at the corresponding pH (donor), as well as an acceptor medium consisting of 2% bovine serum albumin in the corresponding 0.15 M citric acid/phosphate buffer. Sink conditions were maintained by replenishing the acceptor medium at each sampling time point and by the addition of 2% buffered bovine serum albumin. The biomimetic Permeapad® barrier (25 mm, InnoMe GmbH, Espelkamp, Germany) was used to investigate the compounds at a concentration of 50 μM. Constant environmental conditions of 37°C, as well as continuous stirring on the acceptor side at 350 rpm were maintained. Samples were collected from the acceptor medium at 30, 60, 90, 120, 180, and 240 min. An aliquot of 50 µL sample was diluted into the calibration range (0.4 – 100 nM) and precipitated with acetonitrile in a ratio of 1:3 (v/v), immediately vortexed and shaken for 10 min at 800 rpm. Samples were then centrifuged for 5 min at 13.200 xg at room temperature; the supernatant was evaporated under heated nitrogen flow at 50°C and the residue reconstituted in 40/60 DMSO/water (v/v). Quantification was performed using a freshly prepared calibration curve in the same matrix, applying identical sample preparation conditions. Quality controls assured the instrument’s performance during the run. Propranolol served as an internal control with a published P_app_ of 12.03 ± 0.17 × 10^-6^ cm**·**s^-1^ applying Permeapad®.(Majid et al., 2021) For the analysis of compound permeability, the cumulative amount of permeated drug (Q_t_), steady-state flux (J_ss_), and apparent permeability coefficient (P_app_) were calculated using Equation 2 - 4.

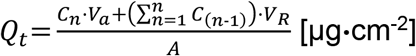

**Equation 2.** Calculation of cumulative amount of permeated drug (Q_t_). C_n_: Drug concentration at time point n, C_n-1_: Drug concentration at previous time point, V_A_: Volume of acceptor chamber, V_R_: Removed volume, A: Permeation area

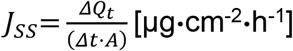

**Equation 3.** Steady-state flux (J_ss_)

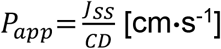

**Equation 4.** Apparent permeability coefficient (P_app_). ΔQ_t_: Difference of Q_t_ between time points, CD: Initial drug concentration, Δt: Time difference, J_ss_: Steady-state flux, A: Permeation area.

## Results

### Molecular Dynamics Simulations of Maurotoxin Reveal Important Interactions

MTx was first docked using the HADDOCK web server. The initial binding mode of MTx was further refined through MD simulations as described in **Methods. Figure 3** summarises the results of the hydrogen bond (H-bond) and cluster analysis conducted with MDAnalysis and GROMACS. The position of MTx does not change significantly throughout the simulations, *i*.*e*., the basic residue Lys23 stays tucked into the pore (**Figure 3A, B**). However, the toxin moves farther across the pore and forms contacts with multiple chains, thereby refining the docking pose (**Figure 3B**). Interacting residues are located around the ion pore and in the turret region (**Figure 3C**). The pose remained stable in five independent simulations (**Figure S5**).

When inspecting the results more closely, important interactions can be identified. One interaction is found between the side chain oxygen of Tyr32 and the backbone oxygens of the channel residues G254, Y253 and D255. This interaction is also recorded between Tyr32 and Q277, which belongs to the turret region of K_Ca_3.1, but only in a smaller fraction of frames. As suggested by the simulations, the three most important interactions are the salt bridge between D239 and the side chain nitrogen atoms of the toxin residues Lys7 or Arg14. The interaction is recorded in more than 60% of simulation frames. The interaction that occurs the most is the H-bond between Lys23 and any of the backbone oxygen atoms of the pore lining residues Y253 and G254. This interaction is formed in more than 70% of simulation frames (**Figure 3D, E**). These results further solidify the importance of the functional dyad and highlight the role of additional positively charged residues that form specific contacts, extending interactions from the pore towards the turret region. Two identified interactions were used as constraints during the virtual screening (**Figure 3F**).

### Docking-based Virtual Screening Campaign

In the first round of docking during the virtual screening, constraints were applied to speed up the calculations. The first H-bond constraint was defined as a H-bond with the oxygen backbone of the ion conductance pore residue Y253 and a H-bond donor atom of the ligand. The second H-bond constraint involves one of the side chain oxygen atoms of D255 and a H-bond donor atom of the ligand. If a molecule cannot fulfil both constraints, it is discarded. This strategy enabled the screening of 48 Mio. molecules in a reasonable amount of time (**Table S1**). After this first rough docking step, the 1 million highest-scoring molecules that fulfil both constraints were selected for redocking. For the more thorough redocking, the constraints were not applied and therefore the molecules were freely placed by the docking algorithm. Next, the generated poses from the free docking step were rescored, this time by applying the constraints to filter out those poses that do not fulfil them. In the rescoring step, all four of GOLD’s scoring functions were used: ChemPLP (Korb et al., 2009), GoldScore (Jones et al., 1995), ChemScore (Verdonk et al., 2003) and ASP (Mooij and Verdonk, 2005). To evaluate the rescoring, a custom Python script was created to further rank the molecules. The ligands were judged according to the following criteria: 1) at least three ligand poses need to fulfil all constraints and they needed to be placed consistently for all scoring functions, 2) docking score, and 3) the poses pass the visual inspection. In total, 3000 poses of 200 molecules were visually inspected to check for correct ligand geometries and interactions beyond the constraints. The aim was to select a structurally diverse set of ligands. Finally, 26 molecules were selected for synthesis (**Figure S2**).

### Validation of Hit Molecule

Initially, the 26 molecules were tested in a fluorescence-based assay in which the membrane potential of K_Ca_3.1 overexpressing HEK293 cells with the voltage-sensitive indicator DiBaC_4_(3) was determined. The channel was activated by increasing the intracellular Ca^2+^ concentration with ionomycin. This resulted in a hyperpolarisation of the membrane potential. We followed a slightly modified procedure as previously published.(Rugi et al., 2025) **1** was the only compound of the initial 26 molecules that showed slight depolarisation which was much weaker than that induced by the known K_Ca_3.1 blocker **12**. The assay may be better suited for detecting potent inhibitors like MTx or **12**, as its sensitivity seems to be limited for weaker binders like initial hits from virtual screening that have not yet undergone lead optimisation. Additionally, the ligands were tested at 10 µM, a concentration that may be insufficient to observe inhibitory effects, particularly considering the solvent-exposed nature of the binding pocket. However, these considerations, combined with the stability of the docking pose of compound **1** in MD simulations (**Figure 4A, Figure S6**), prompted further investigation of its activity through whole-cell patch clamp experiments conducted by ICE Bioscience (Beijing, China). As shown in **Figure 4B**, the compound inhibited K_Ca_3.1 by 21% in a concentration of 250 µM. We therefore decided to expand the structure of this initial hit to obtain inhibitors with improved activity.

**Figure 4.**
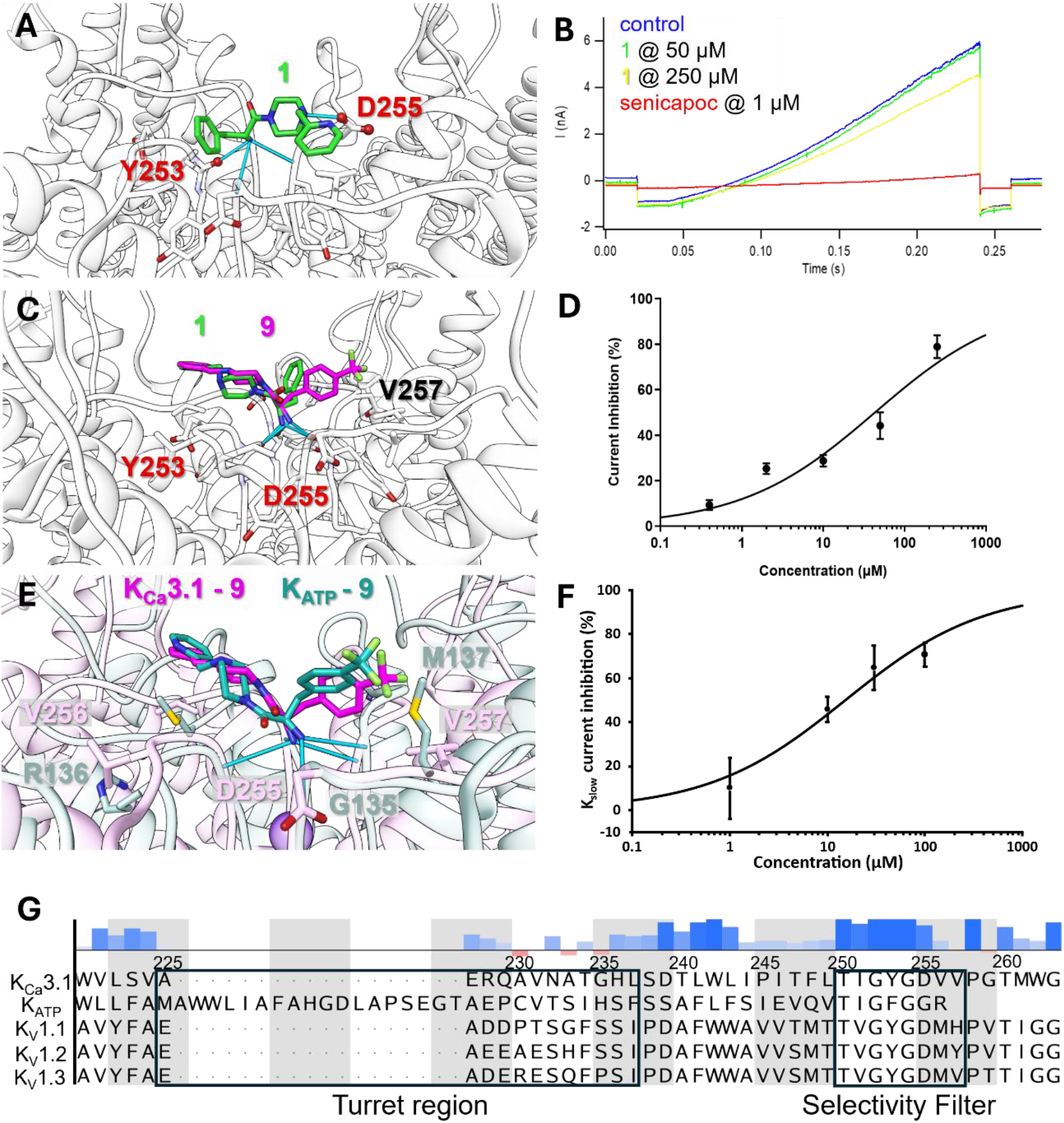
**A**) Docking pose of **1** (green). H-bonds depicted in blue. **B**) Preliminary whole-cell patch clamp results of **1** (n=1). **C**) Docking result of **9** (pink) superimposed on the docking pose of **1** (green). H-bonds depicted in blue. **D**) Inhibition of **9** measured in K_Ca_3.1-overexpressing CHO cells (0.4, 2 and 10 µM: n=2, 50 and 250 µM: n=3). The measured IC_50_ is 43.1 µM. **E**) Docking poses of **9** bound to K_Ca_3.1 (pink) and of **9** bound to K_ATP_ (bluegreen, PDB ID 9KGL). **F**) Inhibition of Ca^2+^-induced K^+^ current by **9** determined in mouse beta cells (1 and 10 µM: n=3; 30 and 100 µM: n=5). The measured IC_50_ is 15.34 µM. **G**) Multiple sequence alignment (MSA) of K_Ca_3.1, K_ATP_ and the voltage-gated ion channels K_V_1.1, K_V_1.2 and K_V_1.3. The numbering depicted above the sequences corresponds to the sequence number of K_Ca_3.1. Furthermore, the sequence similarity is shown in a bar plot: the more positive the bar and the darker blue the more similar the sequences are at this position. Red indicates low similarity and the lower the bar the lower the similarity. The corresponding Uniprot IDs for the sequences of K_Ca_3.1, K_ATP_, K_v_1.1, K_v_1.2, and K_v_1.3 are O15554, Q14654, Q09470, P16389 and P22001, respectively. The webserver CLUSTAL O (v1.2.4) was used for the MSA and the depiction was created with MOE.

### Extended Series Based on Hit Compound 1

The Enamine Real Space was searched via the web interface to identify suitable compounds to expand the initial hit molecule. The structures of matching molecules were downloaded and loaded into Datawarrior (Sander et al., 2015). There, the molecules were clustered and the “Select most diverse selection” option was used to obtain representative molecules of this dataset. The coverage of interesting clusters was visually inspected, and the structures were further filtered based on their visual closeness and Tanimoto score to compound **1**. Moreover, three molecules were selected to investigate whether an aliphatic, primary and freely rotatable amine is necessary for inhibiting K_Ca_3.1 channels. We hypothesised that these three molecules, **4, 7** and **11**, would not efficiently inhibit the channel. Finally, ten molecules were selected, and their respective docking poses are shown in **Figure 5**. The activity of these ten molecules was tested in whole-cell patch clamp assays as described above and the measured activities are listed in **Table 1**. The docking predictions and further computational investigations via MD simulations largely overlap with the experimental results. **5, 9** and **10** significantly improve the activity of the initial hit compound. **9** in particular showed promising activity, with an IC_50_ of 43.1 µM in duplicate whole-cell patch clamp recordings (**Figure 4D**).

**Figure 5.**
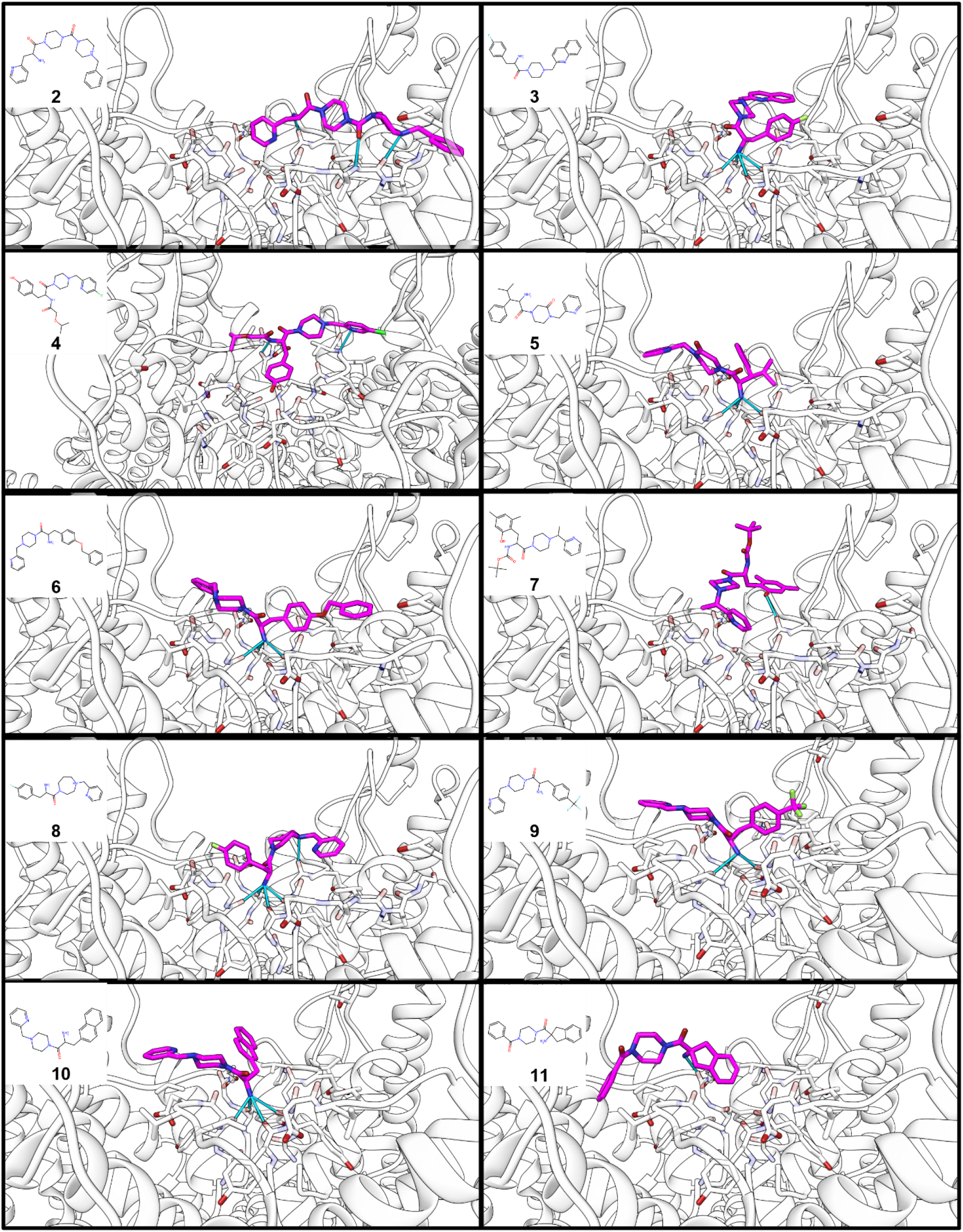
Docking poses of extended hit series (**2-11**).

**Table 1.** Results of the CHO patch clamp experiment for the initial hit (1: n=1) and the compounds of the extension series (2-8, 10, 11: n=1; 9: n=3).

| Compound | Inhibition 50 μM | SE | Inhibition 250 μM | SE |
| --- | --- | --- | --- | --- |
| **1** | 3.24% |  | 23.16% |  |
| **2** | 8.03% |  | 12.65% |  |
| **3** | 10.83% |  | 42.86% |  |
| **4** | 0.36% |  | 42.45% |  |
| **5** | 31.82% |  | 77.09% |  |
| **6** | 8.20% |  | 42.15% |  |
| **7** | 9.17% |  | 34.05% |  |
| **8** | 6.65% |  | 24.42% |  |
| **9** | 44.23% | 5.88% | 78.92% | 5.00% |
| **10** | 41.58% |  | 74.54% |  |
| | Inhibition 30 μM | | Inhibition 100 μM | |
| **11** | -4.05% |  | -2.52% |  |

The molecules **3, 5, 6, 8, 9** and **10** have in common that they all feature a primary amine that is inserted into the ion conductance pore in their respective docking poses. This orientation in the binding site is also stable in the MD simulation of these compounds (**Figure S7**). The primary amine thereby mimics Lys23 of MTx and is blocking ion flux.

As **9** showed the most promising *in vitro* activity, further manual patch clamp assays, performed on murine pancreatic beta cells, were conducted (**Figure 4F**). The K_slow_ current mediated by K_Ca_3.1 and K_ATP_ channels was measured (**Figure S8**).(Düfer et al., 2009) In these experiments, **9** showed a higher activity compared with the patch clamp evaluation in K_Ca_3.1 overexpressing CHO cells. Docking results indicate that **9** can bind to a similar extracellular side in K_ATP_ channels than in K_Ca_3.1 (**Figure 4E**). Additionally, we compared the sequences of K_Ca_3.1 and voltage-gated potassium channels to illustrate the high similarity of the pore region including the selectivity filter and the lower similarity of the turret region (**Figure 4G**).

### Investigation of in vitro Permeability

To further characterise hit compound **1** and the improved inhibitor **9**, their logD values and their permeability were determined in comparison to the intracellularly acting ion channel inhibitor **12** (reference compound senicapoc). Compound **12** showed a logD of 3.18 (mean of n=3) at pH 7.4 and remained pH-independent across the tested pH range (**Table S3**). Its apparent permeability (P_app_) was 2.9 × 10^-6^ cm**·**s^-1^ (mean of n=10) (**Table 2**), consistent with efficient membrane penetration and intracellular target engagement. The novel hit series compounds **1** and **9** exhibited lower lipophilicity with logD_7.4_ of 0.43 and 1.60 (mean of n=3), decreasing to ≤ −2.50 under acidic conditions (pH 2.2). This pronounced pH dependence was reflected in their permeability profile (**Table S3**). While P_app_ values at pH 7.4 indicate good membrane permeability (**Table 2**), permeability decreases markedly at lower pH, reaching 2.2 × 10^−6^ cm**·**s^-1^ and 2.8 × 10^−6^ cm**·**s^-1^ for **1** and **9** at pH 5.0 and 0.3 × 10^−6^ cm**·**s^-1^ and 0.5 × 10^−6^ cm**·**s^-1^ at pH 2.2, respectively. Overall, the reduced lipophilicity and strong pH-dependent permeability distinguish the new compounds **1** and **9** from the intracellular reference inhibitor **12**.

**Table 2.** Apparent permeability coefficient (P_app_) in cm·s^−1^ at different pH values. Mean ± standard error of the depicted n-numbers is shown.

| Compound | $P_{app} (\text{cm} \cdot \text{s}^{-1})$<br>pH 7.4 | n | $P_{app} (\text{cm} \cdot \text{s}^{-1})$<br>pH 5.0 | n | $P_{app} (\text{cm} \cdot \text{s}^{-1})$<br>pH 2.2 | n |
| --- | --- | --- | --- | --- | --- | --- |
| **1** | $11.09 \pm 0.67$<br>$\times 10^{-6}$ | 3 | $2.19 \pm 0.13$<br>$\times 10^{-6}$ | 3 | $0.25 \pm 0.07$<br>$\times 10^{-6}$ | 3 |
| **9** | $8.65 \pm 0.16 \times 10^{-6}$ | 3 | $2.83 \pm 0.08 \times 10^{-6}$ | 3 | $0.46 \pm 0.15 \times 10^{-6}$ | 3 |
| **12** | $3.82 \pm 0.31 \times 10^{-6}$ | 6 | $2.13 \pm 0.06 \times 10^{-6}$ | 2 | $2.71 \pm 0.07 \times 10^{-6}$ | 2 |

## Discussion

In this study, we set out to identify novel extracellular inhibitors of plasma-membrane K_Ca_3.1 channels by combining *in silico* and *in vitro* methods. Therefore, the binding mode of MTx to this channel was studied in detail and important interactions were used as constraints for the docking step. This led to the identification of the initial hit **1**. To further verify the activity of **1**, molecules extending the structure of this initial hit were purchased and tested via whole-cell patch clamp assays.

Except for **11**, all molecules from the extended series can inhibit K_Ca_3.1 channels to some extent, even those that were included as negative controls (**4** and **7**). This might be explained by a different binding mode than suggested here, as the MD simulations indicate that both molecules rapidly unbind from the channel surface. **2** has a primary amine, its docking pose and MD simulation suggest however that it cannot adopt a similar pose as **1**, which explains it comparatively low activity (**Figure S5A**). The flexibility of the primary amine of **11** is restricted and consistent with the computational prediction, no inhibition of the channel was detected. This supports the idea, that in the case of these novel inhibitors, a primary and freely rotatable amine is necessary for inhibition of the K_Ca_3.1 channel to mimic the lysin residue of MTx. The three molecules with improved inhibitory activity compared to **1, 5, 9** and **10**, extend the structure of **1** by adding hydrophobic groups. In the case of **5**, this group is an isopropyl group, while **9** has an additional ortho-trifluoromethyl group, and **10** features a naphthalene moiety instead of a benzene. As suggested by the docking poses, these additional groups enable more hydrophobic interactions, especially with the side chain of V257. **5** is the most stable compound in the MD simulation, while **9** is stable over the whole trajectory in two out of three simulations. The binding mode of **10** is stable in all three simulation repeats.

As the initial patch clamp results indicated that **9** was the most active molecule this compound was further investigated by patch clamp assays performed on murine pancreatic beta cells (**Figure 4F**). These cells not only express K_Ca_3.1 but, amongst others, also ATP- (K_ATP_) and voltage-gated ion channels. In these cells, the main channels contributing to the Ca^2+^-activated K_slow_-current measured in the standard whole-cell configuration are K_Ca_3.1, K_ATP_ and – mediating the Ca^2+^-influx evoked by the depolarizing protocol – voltage-gated calcium channels.(Düfer et al., 2009) The apparent increase in activity in this assay can be explained by the fact that **9** potentially binds to a similar binding site on the K_ATP_ channel (**Figure 4E**). Thus, the selectivity of **9** needs to be improved in the future to optimise its inhibitory potential.

Due to the high similarity of potassium channels in the region surrounding the ion pore, a higher off-target activity was to be expected. However, with this starting point at hand we believe an optimisation into a highly selective extracellular blocker will be possible. Selectivity can presumably be achieved by focusing on interactions in the turret region, which is a loop proceeding the pore helix and influencing the binding of toxins (**Figure 4G**). This domain is dissimilar in K_v_ channels compared with the region surrounding the ion pore, like others also pointed out.(Rauer et al., 2000) In K_v_ channels, this region is more polar and there are more negatively charged residue, for example Q229 of K_Ca_3.1 corresponds to E351(K_v_1.1), E353 (K_v_1.2) and D423 (K_v_1.3). Therefore, introducing carboxylic acid moieties is a promising strategy to mitigate activity on K_v_ channels. K_ATP_ channels on the other hand have an elongated turret region and therefore more spacious molecules might lead to reduced affinity to this channel.

Another area of improvement is the existing membrane permeability of the identified small molecule inhibitor. This permeability reduces the effective concentration of the extracellular inhibitor and leaves room for further optimisation. As the permeability and logD measurements suggest, these properties of **9** are highly dependent on the pH of the surrounding environment. Incorporation of a quaternary amine moiety, for example, represents a potential strategy to improve the inhibitors. Such a modification is expected to further restrict membrane permeability, as indicated by the low P_app_ values at pH 2.2 at which the compounds are highly ionised.

Previous efforts have identified the cyclised dequalinium derivative UCL1684, which was recently confirmed to block K_Ca_2.2 from the extracellular side.(Rosa et al., 1998, 2000; Nam et al., 2025) This blocker lacks activity on K_Ca_3.1. However, other dequalinium derivatives were shown to be able to block K_Ca_3.1. For example, UCL1144 is 15 times more effective in blocking K_Ca_3.1 than K_Ca_2.2, but blocks both channels in the micromolar range, with 1.22 µM and 18 µM, respectively. UCL1144 contains two 4-hydroxyquinoline moieties connected via a linear decane chain.(Malik-Hall et al., 2000) It therefore lacks the cyclised structure and quaternary amine of UCL1684 and it can be assumed that it is a rather unselective hydrophobic binder which does not bind to the extracellular side of K_Ca_3.1. Moreover, the molecules from the UCL series have a low drug likeness. Therefore, we claim that **9** is the first drug-like, small-molecule extracellular inhibitor of K_Ca_3.1 with the huge potential to be developed into a selective K_Ca_3.1 channel blocker.

## Conclusions

This work presents the identification of **9**, a new small molecular drug-like inhibitor of K_Ca_3.1 that blocks the channel from the extracellular side. Due to the increased speed of docking calculations with constraints, it was possible to virtually screen the Molport database, which contains over 48 million purchasable molecules. This process led to the identification of novel compounds, which were further examined using MD simulations. Promising molecules were purchased and tested *in vitro* to confirm their ability to inhibit K_Ca_3.1 channels. Future optimisation strategies will prioritise selective engagement of the extracellular turret region and tighter control of membrane permeability, thereby minimising the potential loss of activity associated with compound permeation to the intracellular compartment.

Moreover, we further examined the binding mode of the known toxin inhibitor MTx and emphasises the importance of Lys23 for blocking K_Ca_3.1. Important residues to guide the modification of MTx to enhance its K_Ca_3.1 blocking ability or facilitate its use as a radio-labelled ligand were identified. Furthermore, these findings are valuable for designing small peptides with greater specificity for K_Ca_3.1 channels than the toxin studied here.

## Author Contributions

J.M. and O.K. conceived and conceptualized the study. J.M. carried out the computational work, performed the analyses, prepared the initial manuscript draft and carried out in-vitro experiments. M.G. supported data acquisition. J.H. and T.G. carried out the in-vitro experiments and performed associated data analyses. E.E.B contributed to in-vitro data acquisition. B.B.B., M.D., A.S. and O.K. contributed to experimental design and provided critical input during manuscript preparation. O.K. was responsible for funding acquisition and supervision. All authors reviewed, edited and approved the final manuscript.

## Acknowledgements

Patch clamp of CHO cells were conducted by ICE Bioscience and molecules were bought from Enamine and through Molport.

This work was supported by the Research Training Group “Chemical biology of ion channels (Chembion)” funded by the *Deutsche Forschungsgemeinschaft* (DFG), which is gratefully acknowledged. O.K. was furthermore funded through a Heisenberg-Professorship (DFG; KO 4689/5–2). J.M. and O.K. gratefully acknowledge the scientific support and HPC resources provided by the Erlangen National High Performance Computing Center (NHR@FAU) of the Friedrich-Alexander-Universität Erlangen-Nürnberg (FAU) under the NHR project k101ee NHR funding is provided by federal and Bavarian state authorities. NHR@FAU hardware is partially funded by the DFG–440719683.

## Data Availability Statement

The modelled toxin-K_Ca_3.1 binding poses are available from the University of Münster datastore: DOI: 10.17879/450fg-qch56.

Further data supporting the findings of this study are available from the corresponding author upon reasonable request.

