## Supporting Information for "Discovery of the first small-molecule extracellular inhibitor of K_Ca_3.1"

Homepage: www.kochlab.org

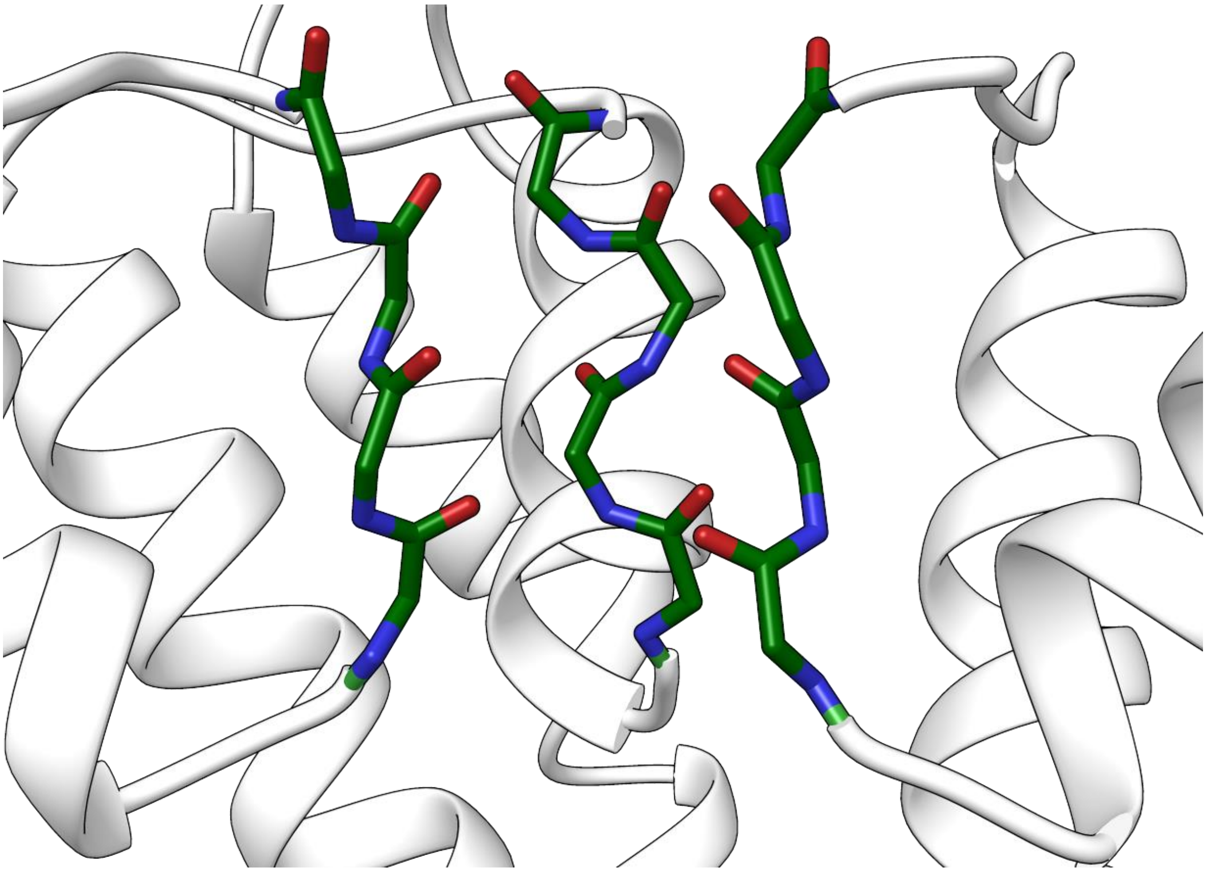

**Figure S1:** Close-up view of the ion conductance pore. Chain D was hidden to enhance visibility. Backbone atoms are displayed as sticks and other protein residues are shown as ribbons. The backbone carbons, nitrogens and oxygens are coloured dark green, blue and red, respectively. The backbone oxygens lose their ordered state if no $K^+$ ions are modelled in the pore. This distortion was also observed for other toxins by others.

**Table S1:** Docking of Molport Database on HPC Cluster Palma II in Münster

| | First run | Redocking | Rescoring |
|---|---|---|---|
| **N° of Ligands** | 48.5 Mio. | 1 Mio. | 1 Mio. |
| **Scoring Function** | ChemPLP | ChemPLP | all[a] |
| **Constraints** | Yes | No | Yes |
| **Poses** | 10[b] | 15 | 15 |
| **GA setting** | 0.3 | 1.0 | 1.0 |
| **Compute Time [h]** | 26,561 | 7,109 | - |
| **Time/Ligand [s]** | 2.1 | 25.6 | - |
| **Real Time** | ~ 2 weeks | 1 day | 4 days |

[a]All four scoring functions implemented in GOLD: ChemPLP(Korb et al., 2009), GoldScore(Jones et al., 1995), ChemScore(Verdonk et al., 2003), Astex Statistical Potential (ASP)(Mooij and Verdonk, 2005) [b]Early termination enabled

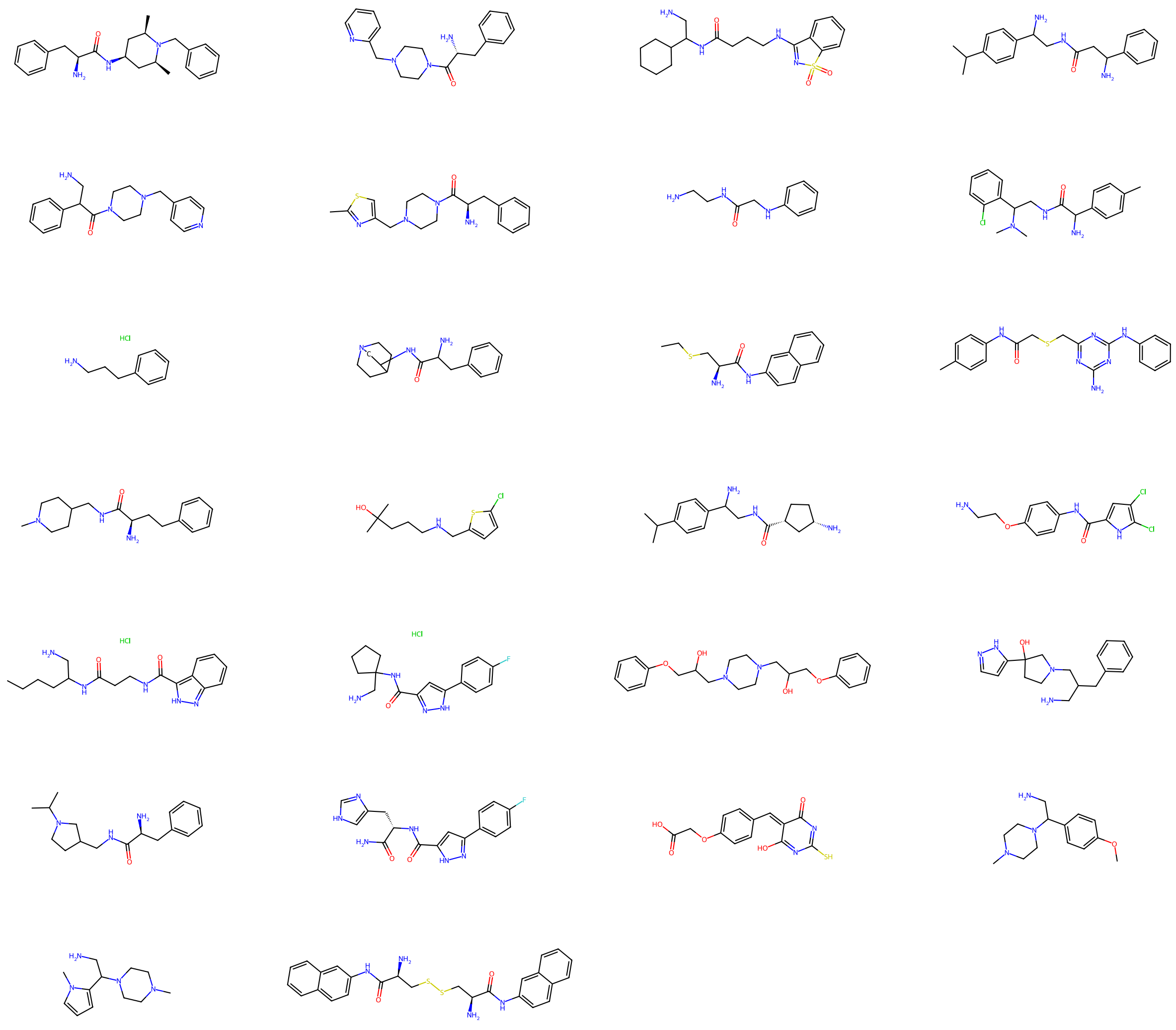

**Figure S2**: Structures of the 26 compounds from the first selection.

**Table S2**: Detailed description and overview about all conducted MD simulations.

| System | Repeat | Simulation Time [ns] | N° of atoms | Box size [Å$^3$] |
|---|---|---|---|---|
| $K_{Ca}$3.1 - MTx | 1 | 513 | 637,569 | 177x177x236 |
| | 2 | 576 | 637,569 | 177x177x236 |
| | 3 | 681 | 637,569 | 177x177x236 |
| | 4 | 502 | 637,569 | 177x177x236 |
| | 5 | 511 | 637,569 | 177x177x236 |
| $K_{Ca}$3.1 - **1** | 1 | 726 | 652,000 | 155x155x267 |
| | 2 | 785 | 652,000 | 155x155x267 |
| | 3 | 727 | 652,000 | 155x155x267 |
| $K_{Ca}$3.1 - **2** | 1 | 500 | 315,487 | 144x144x147 |
| | 2 | 500 | 315,487 | 144x144x147 |
| | 3 | 500 | 315,487 | 144x144x147 |
| $K_{Ca}$3.1 - **3** | 1 | 500 | 315,508 | 144x144x147 |
| | 2 | 500 | 315,508 | 144x144x147 |
| | 3 | 500 | 315,508 | 144x144x147 |
| $K_{Ca}$3.1 - **4** | 1 | 500 | 315,427 | 145x145x146 |
| | 2 | 500 | 315,427 | 145x145x146 |
| | 3 | 500 | 315,427 | 145x145x146 |
| $K_{Ca}$3.1 - **5** | 1 | 500 | 315,381 | 144x144x147 |
| | 2 | 500 | 315,381 | 144x144x147 |
| | 3 | 500 | 315,381 | 144x144x147 |
| $K_{Ca}$3.1 - **6** | 1 | 500 | 315,638 | 145x145x146 |
| | 2 | 500 | 315,638 | 145x145x146 |
| | 3 | 500 | 315,638 | 145x145x146 |
| $K_{Ca}$3.1 - **7** | 1 | 500 | 315,151 | 144x144x148 |
| | 2 | 500 | 315,151 | 144x144x148 |
| | 3 | 500 | 315,151 | 144x144x148 |
| $K_{Ca}$3.1 - **8** | 1 | 500 | 315,575 | 144x144x148 |
| | 2 | 500 | 315,575 | 144x144x148 |
| | 3 | 500 | 315,575 | 144x144x148 |
| $K_{Ca}$3.1 - **9** | 1 | 500 | 315,164 | 144x144x148 |
| | 2 | 500 | 315,164 | 144x144x148 |
| | 3 | 500 | 315,164 | 144x144x148 |
| $K_{Ca}$3.1 - **10** | 1 | 500 | 315,337 | 144x144x146 |
| | 2 | 500 | 315,337 | 144x144x146 |
| | 3 | 500 | 315,337 | 144x144x146 |
| $K_{Ca}$3.1 - **11** | 1 | 500 | 315,470 | 144x144x147 |
| | 2 | 500 | 315,470 | 144x144x147 |
| | 3 | 500 | 315,470 | 144x144x147 |

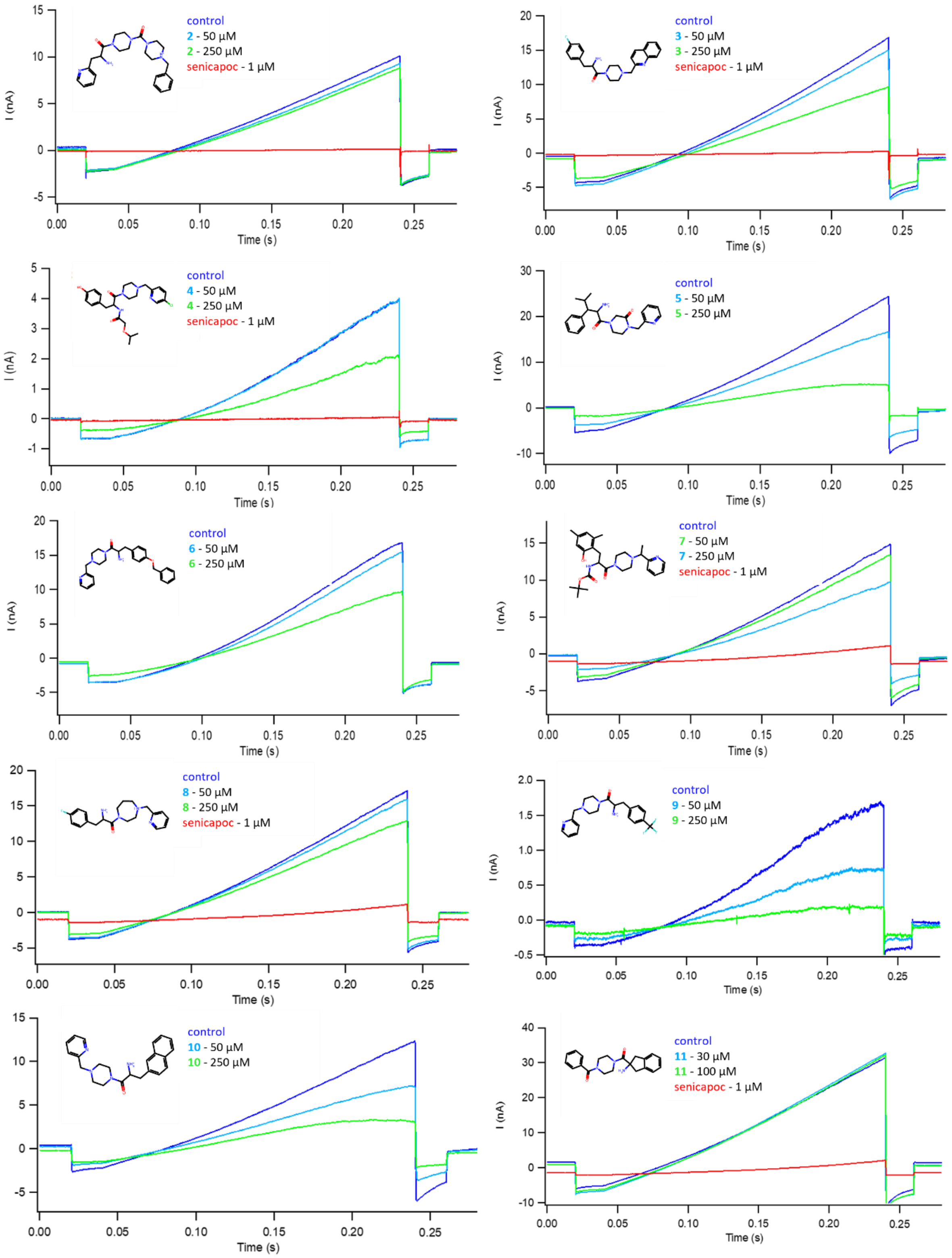


**Figure S3**: Results of whole-cell patch clamp assay. Individual curves of **2-11** (n=1).

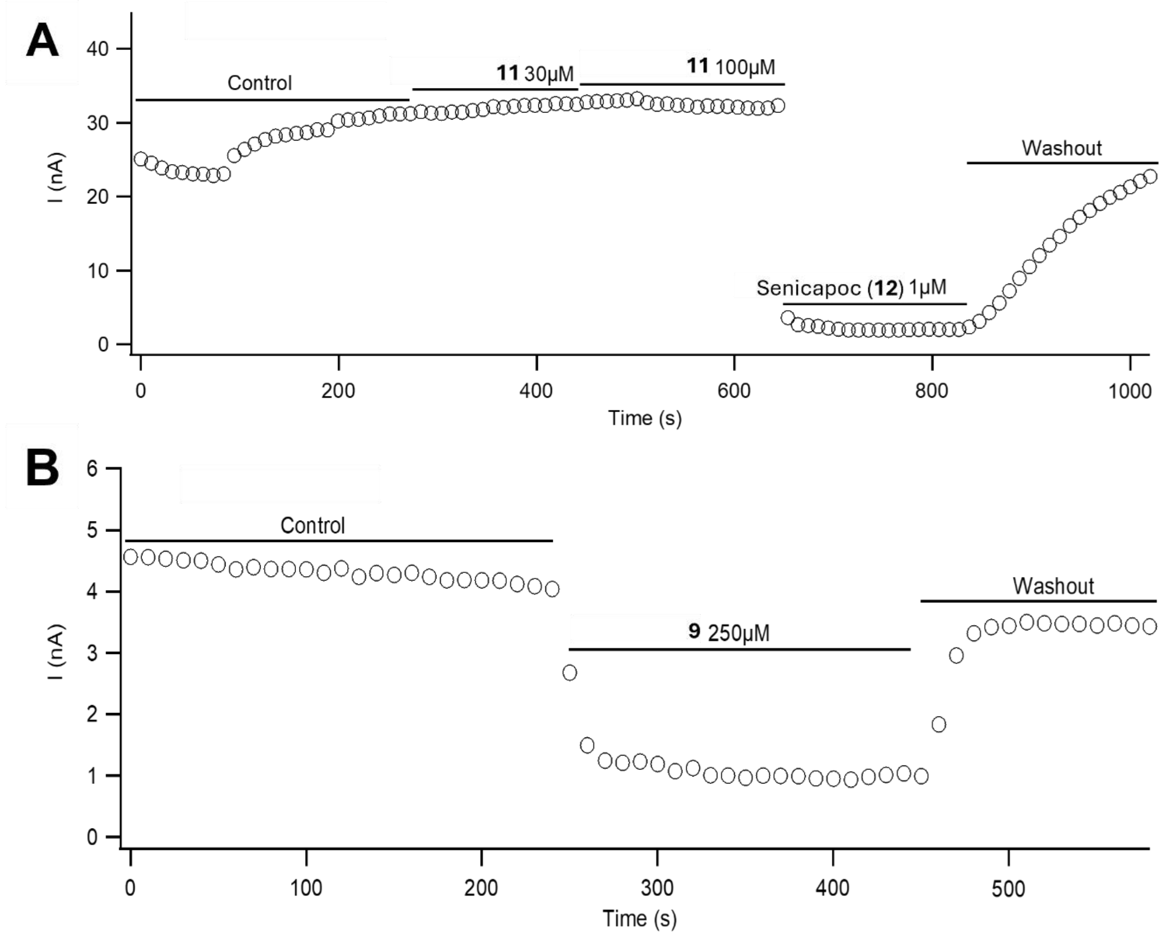


**Figure S4**: Patch clamp traces from the testing of the compounds on $K_{Ca}3.1$ overexpressing CHO cells. Exemplary results for **A**) **11** (n=1) and **B**) **9**. Depicted is a single experiment of **9** (n=1). However, **9** was measured in total three times at 250 µM (n=3).

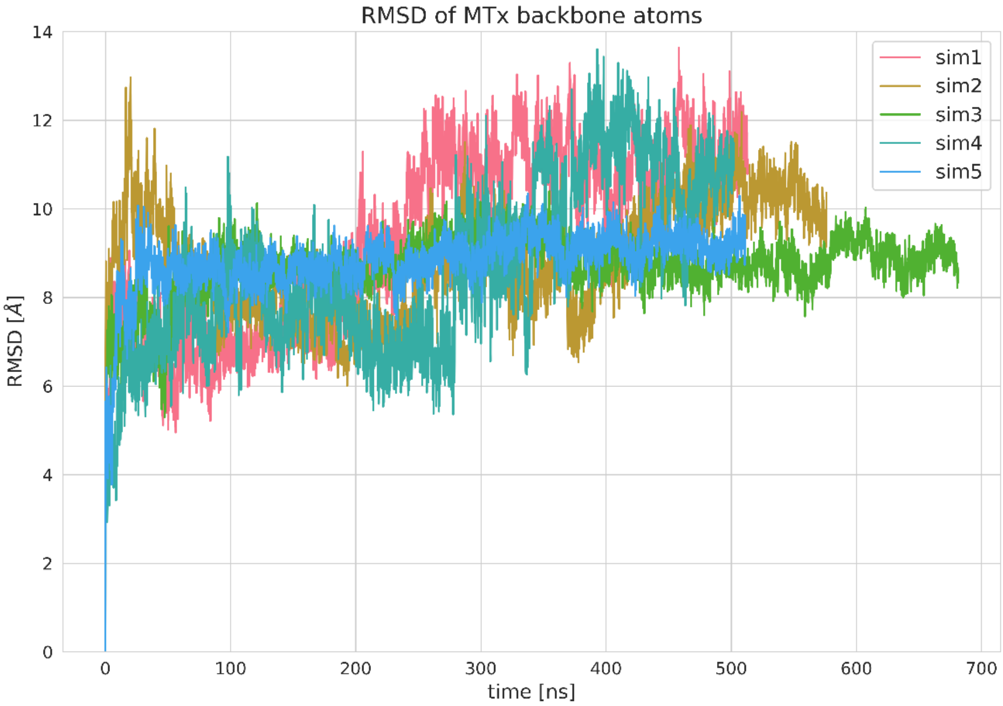


**Figure S5**: RMSD of Maurotoxin backbone atoms.

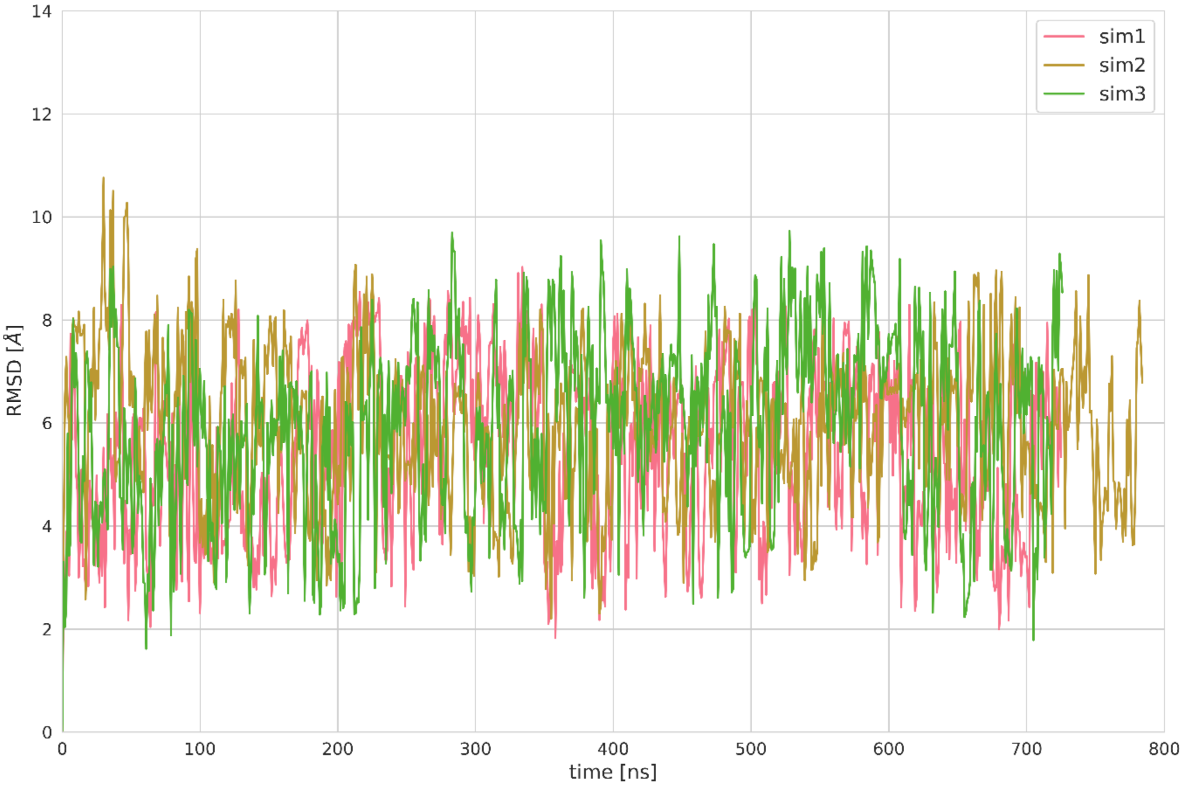


**Figure S6**: RMSD of **1** heavy atoms.

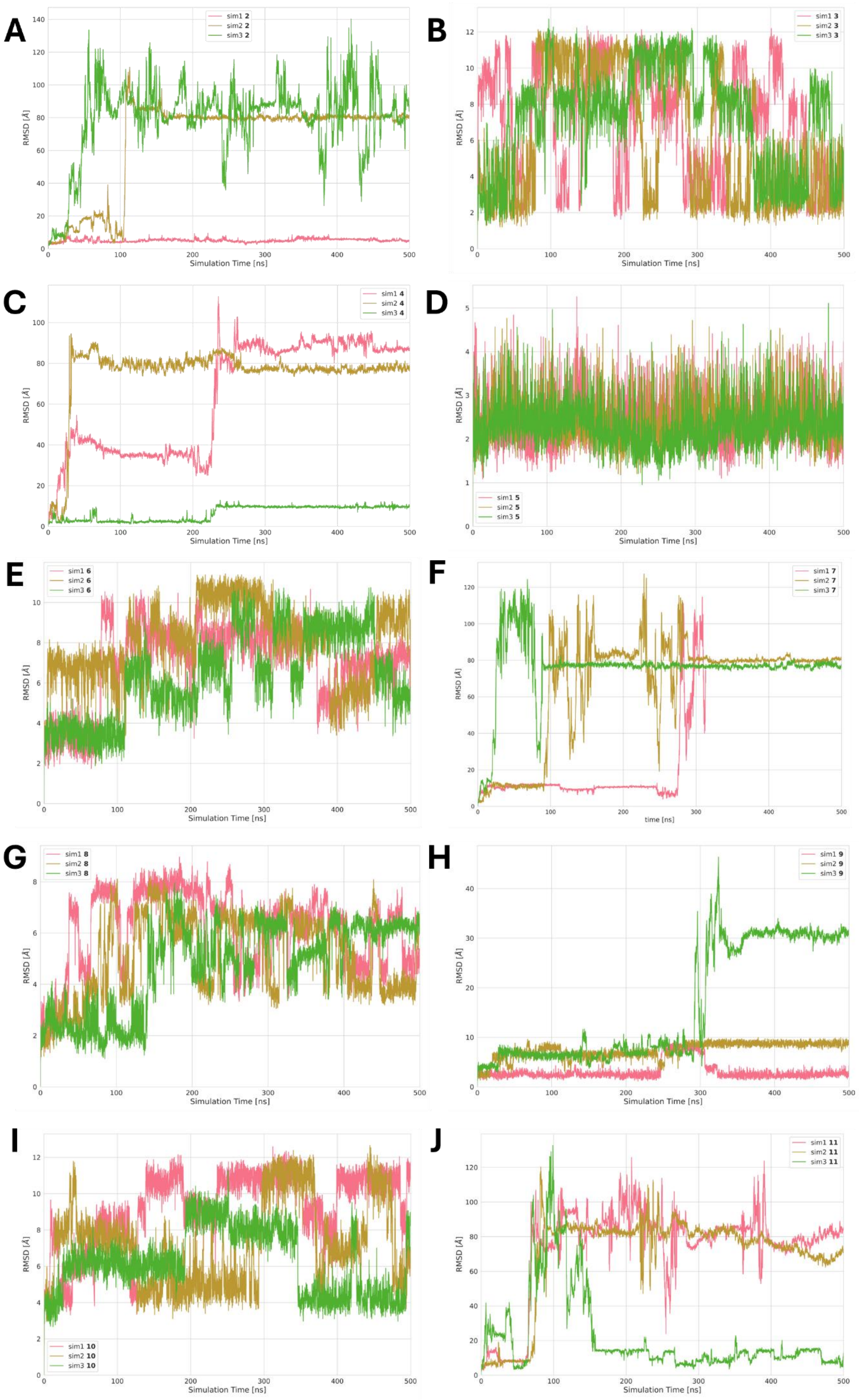


**Figure S7**: RMSD of **2-11** heavy atoms (**A-J**).

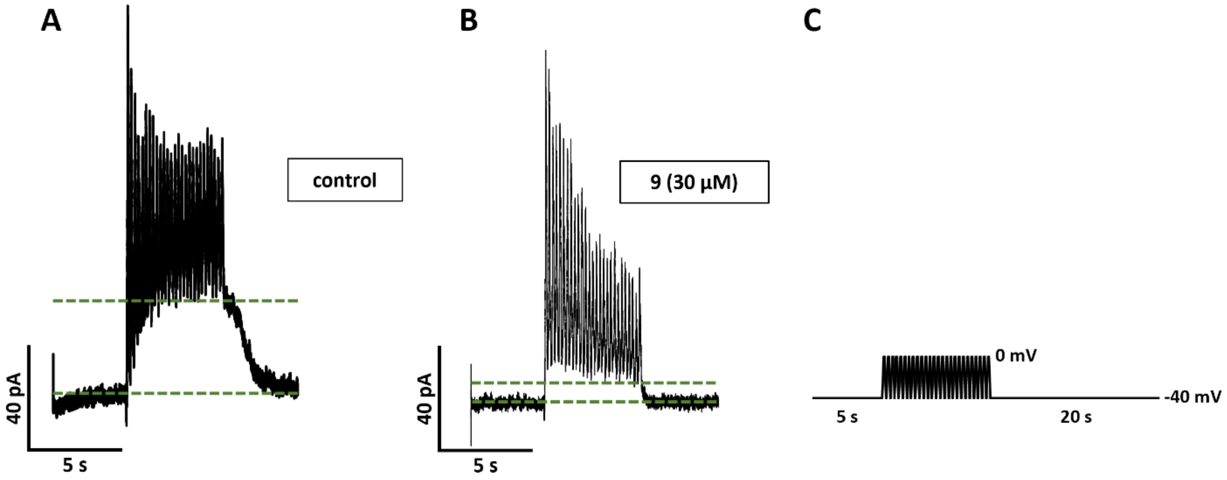


**Figure S8**: Exemplary traces from the testing of **9** on murine beta cells. Whole-cell currents were evoked by a train of voltage ramps (-40 to 0 mV, 26x, **C**) before **A**) and in the presence **B**) of the compound (n=5). Green lines in A and B: amplitude of $K_{slow}$ current.

**Table S3**: Determined logD values. Data are presented as mean ± SD (n = 3).

| Compound | $logD_{7.4}$ | $logD_{5.0}$ | $logD_{2.2}$ |
|---|---|---|---|
| **1** | 0.43 ± 0.02 | -1.85 ± 0.07 | -2.50 ± 0.22 |
| **9** | 1.60 ± 0.08 | -0.66 ± 0.02 | -2.72 ± 0.18 |
| **12** | 3.29 ± 0.08 | 3.27 ± 0.15 | 3.21 ± 0.04 |
